# Comparison of two multi-trait association testing methods and sequence-based fine mapping of six QTL in Swiss Large White pigs

**DOI:** 10.1101/2022.12.13.520268

**Authors:** A. Nosková, A. Mehrotra, N.K. Kadri, A. Lloret-Villas, S. Neuenschwander, A. Hofer, H. Pausch

**Affiliations:** ETH Zürich, Universitätstrasse 2, 8092, Zürich, Switzerland; ETH Zürich, Tannenstrasse 1, 8092, Zürich, Switzerland; SUISAG, Allmend 10, 6204 Sempach, Switzerland

**Keywords:** Genome-wide association study, Multivariate analyses, Meta-analyses, Pleiotropy, Imputation

## Abstract

**Background:** Genetic correlations between complex traits suggest that pleiotropic variants contribute to trait variation. Genome-wide association studies (GWAS) aim to uncover the genetic underpinnings of traits. Multivariate association testing and the meta-analysis of summary statistics from single-trait GWAS enable detecting variants associated with multiple phenotypes. In this study, we used array-derived genotypes and phenotypes for 24 reproduction, production, and conformation traits to explore differences between the two methods and used imputed sequence variant genotypes to fine-map six quantitative trait loci (QTL).

**Results:** We considered genotypes at 44,733 SNPs for 5,753 pigs from the Swiss Large White breed that had deregressed breeding values for 24 traits. Single-trait association analyses revealed eleven QTL that affected 15 traits. Multi-trait association testing and the meta-analysis of the single-trait GWAS revealed between 3 and 6 QTL, respectively, in three groups of traits. The multi-trait methods revealed three loci that were not detected in the single-trait GWAS. Four QTL that were identified in the single-trait GWAS, remained undetected in the multi-trait analyses. To pinpoint candidate causal variants for the QTL, we imputed the array-derived genotypes to the sequence level using a sequenced reference panel consisting of 421 pigs. This approach provided genotypes at 16 million imputed sequence variants with a mean accuracy of imputation of 0.94. The fine-mapping of six QTL with imputed sequence variant genotypes revealed four previously proposed causal mutations among the top variants.

**Conclusions:** Our findings in a medium-size cohort of pigs suggest that multivariate association testing and the meta-analysis of summary statistics from single-trait GWAS provide very similar results. Although multi-trait association methods provide a useful overview of pleiotropic loci segregating in mapping populations, the investigation of single-trait association studies is still advised, as multi-trait methods may miss QTL that are uncovered in single-trait GWAS.

## Background

Genome-wide association studies (GWAS) combine genotype and phenotype information to identify trait-associated variants. Genotypes at polymorphic loci are tested for association with phenotypes to determine their impact on traits of interest. Multi-trait GWAS can increase the statistical power over single-trait GWAS because they exploit cross-phenotype associations at pleiotropic loci [1–3].

Several methods have been developed to detect pleiotropic variants. These methods can be divided into two groups based on their underlying statistical framework [4, 5]. First, multivariate methods jointly model all traits of interest. This group of methods requires that all individuals included in the study have phenotypic records for all traits analysed, although there are exceptions (e.g., single step GWAS [6], imputation of phenotypes [7]). These methods exploit the genetic covariance between traits, thereby increasing statistical power over their univariate counterparts [5, 8, 9], unless all traits are highly correlated [9, 10]. Second, the meta-analysis of summary statistics enables to combine results from single trait GWAS, which means that the analyses can be carried out with different sets of individuals for each trait [1, 11–13].

The power to detect trait-associated variants increases as the marker density increases [14]. Low-pass sequencing is a cost-effective approach to provide high marker density [15–18]. The imputation from medium-density arrays to the whole-genome sequence level using a sequenced reference panel is another approach to provide sequence variant genotypes for large cohorts. The imputation of sequence variant genotypes is typically performed in two steps [19–21]. Imputation from medium density genotypes to the whole genome sequence level has been explored when high-density array-derived genotypes were not available [22–25].

Only few genome-wide association studies have been conducted in the Swiss Large White (SLW) population. Becker et al. [26] performed association tests between 26 complex traits and 60K SNPs genotyped in 192 breeding boars. This effort revealed only 4 QTL likely because the sample size was too small. Large-scale association testing had been conducted in other pig breeds (e.g., [27, 28, 29]). Fat deposition and weight gain-related traits have been considered frequently in these GWAS as they are economically relevant and highly heritable. Previous GWAS led to tens of proposed candidate genes affecting these traits, including *MC4R, BMP2, IGF2*, and *CCND2* [19, 30–34].

In this paper, we compare single-trait, multivariate and meta-GWAS in 5,753 genotyped pigs from a Swiss breed to investigate the genetic architecture of 24 traits. We inferred sequence variant genotypes from a sequenced reference panel to identify candidate causal variants for six pleiotropic QTL.

## Results

### Single-trait association studies between array genotypes and 24 traits

A detailed description of the 24 traits considered in our study including their grouping into four categories (reproduction, production, conformation, all) is shown in Table 1. Marked pairwise correlations exist between the drEBV of the 24 traits (Additional file 1). The SNP-based heritability estimates of the drEBV (Table 1) were between 0.04 and 0.67.

**Table 1.**
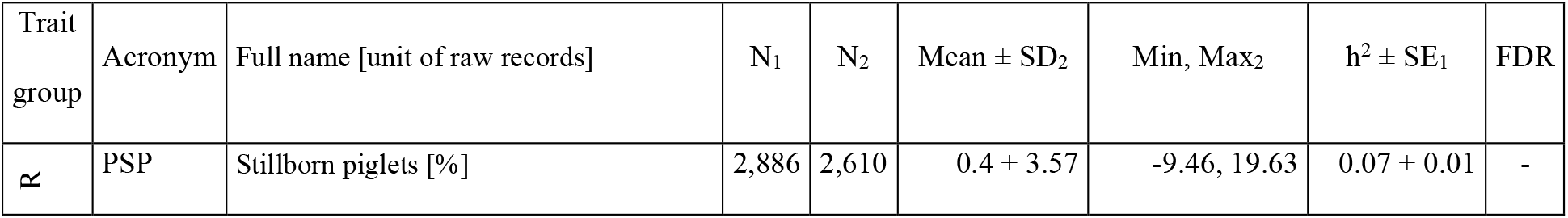
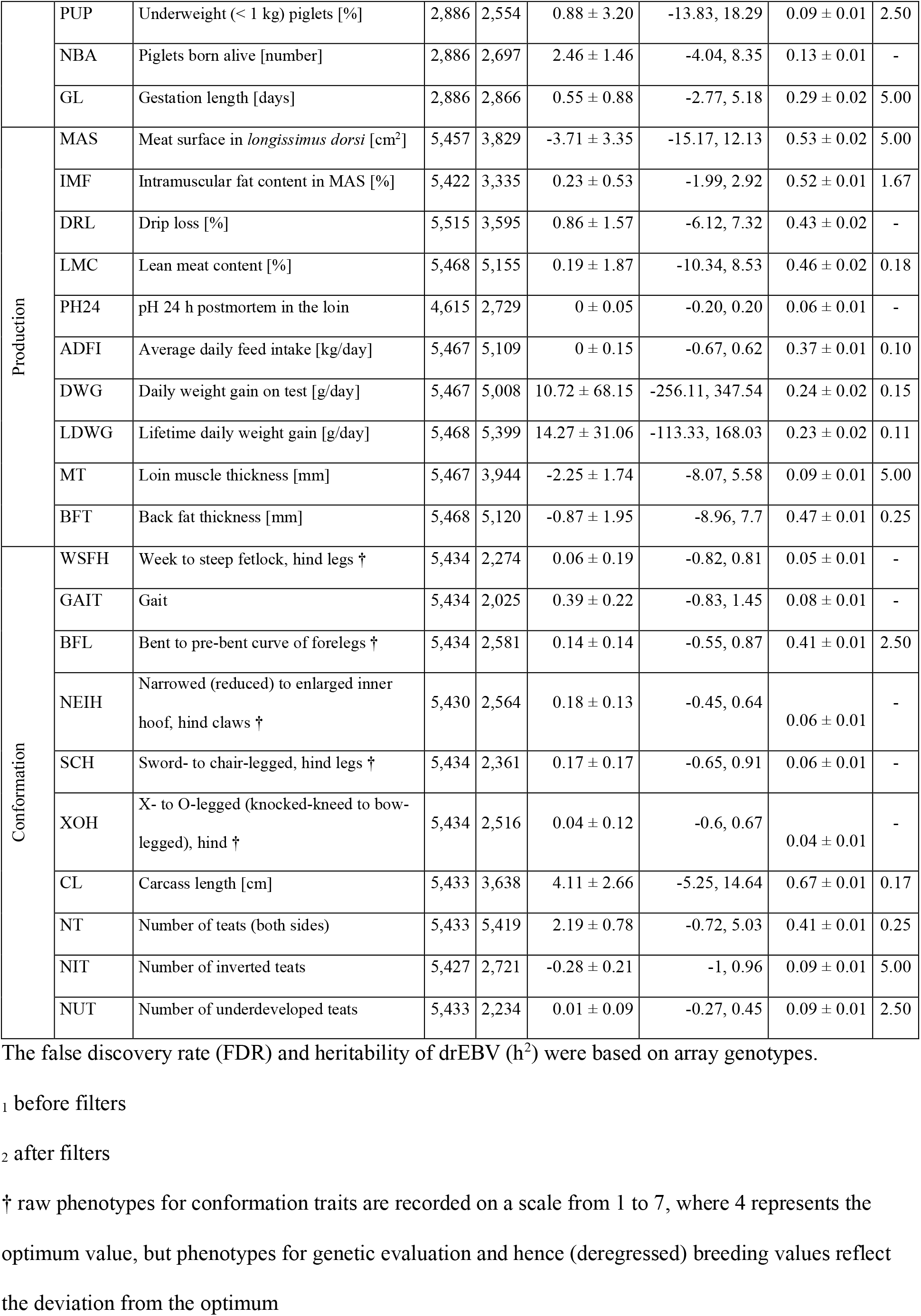
Traits with their abbreviations, full descriptions, corresponding trait group and descriptive statistics of the drEBV.

Mixed model-based single-trait genome-wide association studies (stGWAS) between 40,382 SNPs and deregressed estimated breeding values (drEBV) for 24 traits in 5,753 Swiss Large White (SLW) pigs revealed 237 significantly associated variants (P < 1.24 × 10^−6^). Fifteen out of 24 traits had at least one significantly associated variant (Table 1; Additional file 2; Additional file 3).

The number of variants that exceeded the Bonferroni-corrected significance threshold was between 1 for GL, NIT, MAS and MTF, and 49 for ADFI (Table 1). The inflation factors of the stGWAS were between 0.85 for NT and 1.03 for SCH with an average value of 0.95 ± 0.04 across all 24 stGWAS indicating that population stratification was properly considered.

The 237 associations were detected at 99 unique SNPs located at 11 QTL on SSC1, 5, 11, 15, 17 and 18. The two strongest associations were detected between the number of teats (NT) and a variant on SSC7 (MARC0038565 at 97,652,632 bp, P: 3.35 × 10^−35^), and between lifetime daily weight gain (LDWG) and a variant on SSC1 (ASGA0008077 at 270,968,825 bp, P: 3.28 × 10^−28^). 57 SNPs were significantly associated with more than one trait (two SNPs, ALGA0123414 and ASGA0008077, were associated with six traits) suggesting that pleiotropic effects are present and detectable in our dataset.

### Comparison of multi-trait studies using array genotypes

In order to exploit genetic correlations among the traits to detect pleiotropic loci, we conducted multivariate linear mixed model-based (mtGWAS) association testing and performed a multi-trait meta-analyses of the single-trait GWAS (metaGWAS^1^) for traits within the four trait categories. For unbiased investigation of differences between both methods, we considered between 1,074 and 2,689 individuals with complete phenotypic records for all traits within a trait category (Table 2) for the multi-trait analyses.

**Table 2.** Number of QTL in trait groups revealed by each of the methods.

| Group | Reproduction | Production | Conformation | All |
| --- | --- | --- | --- | --- |
| Number of traits | 4 | 10 | 10 | 24 |
| Number of animals with records for all traits | 2,553 | 1,927 | 2,689 | 1,074 |
| Mean correlation between drEBV (± SD) | $0.14 \pm 0.18$ | $0.32 \pm 0.24$ | $0.17 \pm 0.19$ | $0.10 \pm 0.16$ |
| Mean $h^2$ (± SD) | $0.14 \pm 0.08$ | $0.35 \pm 0.16$ | $0.16 \pm 0.20$ | $0.23 \pm 0.19$ |
| QTL - stGWAS<sup>1*</sup> (in N traits) | 2 (2) | 5 (8) | 4 (5) | 11 (15) |
| QTL - mtGWAS<sup>1*</sup> | 0 | 5 | 3 | 4 |
| QTL - metaGWAS<sup>1*</sup> | 0 | 7 | 3 | 4 |
| QTL - metaGWAS<sup>2*</sup> | 0 | 6 | 3 | 7 |
| QTL - metaGWAS<sup>2</sup> (imputed WGS) | 0 | 6 | 3 | 6 |

<sup>1</sup> performed in pigs with complete records in the trait groups
<sup>2</sup> performed in GWAS conducted in between 2,025 and 5,419 pigs
\* based on array-derived genotypes at 40,382 SNPs

Both methods yielded similar results, but the metaGWAS^1^ revealed more significantly associated variants as well as QTL (Table 2; Figure 1; Additional files 4-6). Across the four trait categories, the metaGWAS^1^ revealed slightly more significant SNPs (Figure 1A) resulting in a 18% smaller FDR than mtGWAS. The metaGWAS^1^ revealed 65 unique variants that were significantly associated with at least one trait category, of which 41 were also detected using mtGWAS. The mtGWAS revealed only associations that were also detected by metaGWAS^1^(Figure 1B). The P-values of lead SNPs were slightly lower (i.e., more significant) in the metaGWAS^1^ than the mtGWAS (Figure 1C).

**Figure 1.**
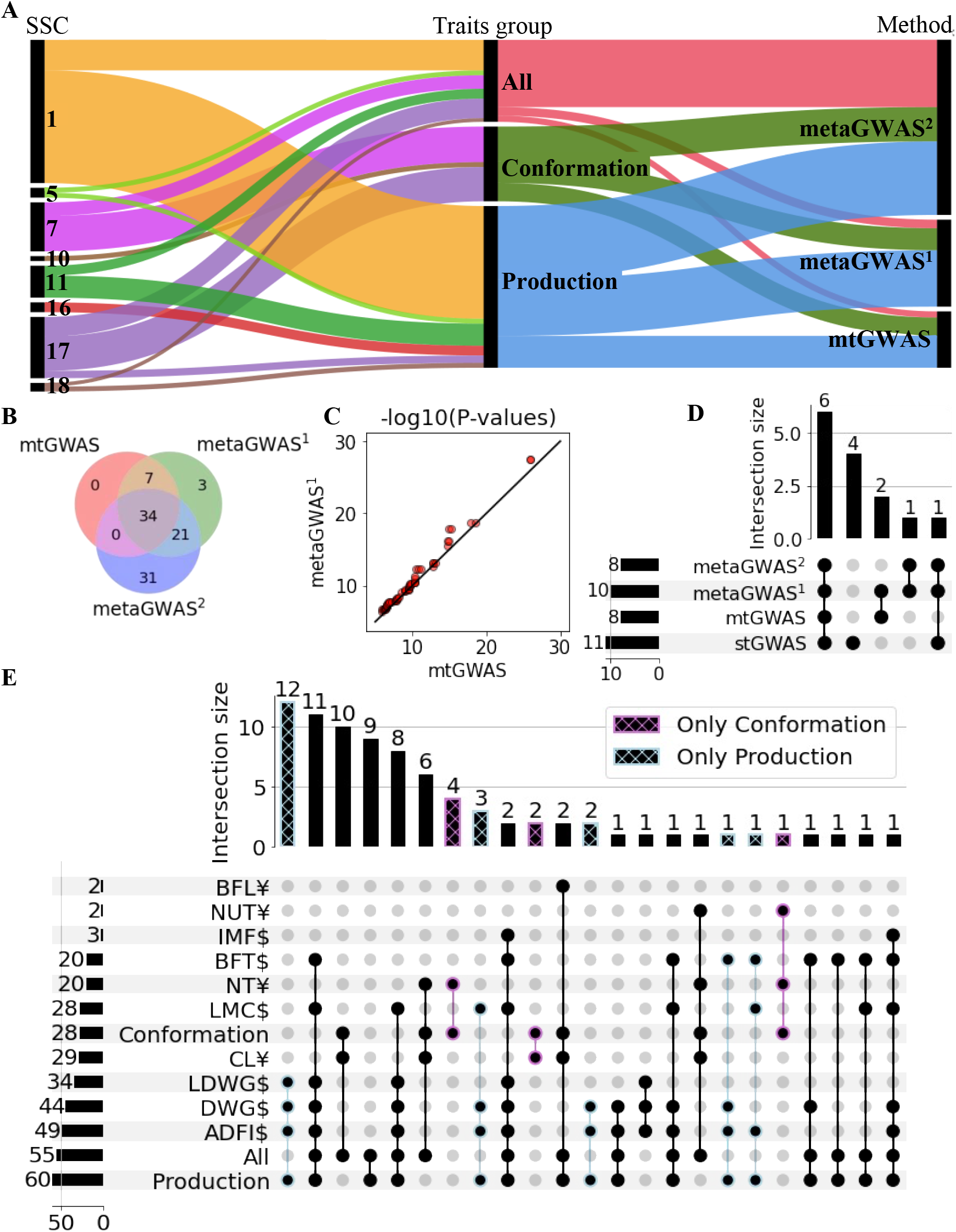
Comparison of variants associated with 24 traits from 3 multi-trait GWAS methods. Multivariate (mtGWAS), meta-analyses with complete dataset (metaGWAS^1^), and meta-analyses including samples with missing trait records (metaGWAS^2^) were based on array-derived genotypes. (A) Proportions of significantly associated variants discovered across chromosomes, groups of traits and multi-trait methods. (B) Overlaps between the associated variants revealed by each of the methods. Sum across all four trait-groups. (C) QQ plot between-log10(P) of variants (N = 41) associated in both mtGWAS and metaGWAS^2^. The line denotes a correlation of 1. (D) QTL detected by different methods across all trait groups. (E) Number of significantly associated pleiotropic variants in the metaGWAS^2^ within groups and stGWAS of individual traits ($ production, ¥ conformation). The single traits are: BFL - Bent to pre-bent curve of forelegs; NUT - Number of underdeveloped teats; IMF - Intramuscular fat content in MAS; BFT - Back fat thickness; NT - Number of teats (both sides); LMC - Lean meat content; CL - Carcass length; LDWG - Lifetime daily weight gain; DWG - Daily weight gain on test; ADFI - Average daily feed intake.

Neither of the multi-trait methods detected significantly associated SNP for the reproduction trait category. For the conformation trait category, mtGWAS and metaGWAS^1^ revealed 15 and 18 associated SNPs, respectively. The associated SNPs defined three QTL on SSC7, 10, and 17. For the production trait category, the mtGWAS and metaGWAS^1^ revealed 26 and 46 associations, respectively. Both methods revealed QTL at SSC1 ~ 270 Mb, SSC16, two QTL at SSC17 ~ 3 Mb and ~ 16 Mb, and SSC18. The metaGWAS^1^ revealed two additional QTL at SSC1 ~ 150 Mb and SSC11. When all 24 traits were combined for 1,074 pigs, the mtGWAS and metaGWAS^1^ revealed only 5 and 7 associated SNPs, respectively. These SNPs spanned four QTL on SSC5, 7, 17, and 18.

Seven QTL detected by mtGWAS and metaGWAS^1^, were also detected by stGWAS (Figure 1D). Both multi-trait methods detected three QTL on SSC11, 16 and 17 ~ 3 Mb that were not detected in the stGWAS. Four QTL detected in the stGWAS were not revealed by either of the multi-trait methods: these were QTL on SSC7, 11, and 15 that were slightly above the Bonferroni-corrected significance threshold for one trait (GL with P = 5.03 × 10^−7^, NIT with P = 1.06 × 10^−6^, PUP with P = 2.66 × 10^−7^, respectively), and one QTL on SSC10 that was associated with MT (P = 5.60 × 10^−7^) and MES (P = 8.44 × 10^−7^). From the seven QTL detected by both multi-trait and single-trait methods, six were associated with more than one trait in stGWAS. For the six pleiotropic QTL, at least one single trait analyses revealed more associated variants, and smaller P-value of the top SNP, than the mtGWAS or metaGWAS^1^(Additional file 7).

Using summary statistics from stGWAS for a multi-trait metaGWAS facilitates including data from animals with partially missing phenotypes. In order to maximize the power to identify trait-associated pleiotropic variants, we reran the metaGWAS using summary statistics from stGWAS with all available animals per trait (Additional file 8), denoted as metaGWAS^2^.

Compared to the previous metaGWAS^1^ that was based on fewer individuals that had phenotypes for all traits, the number of SNPs exceeding the Bonferroni-corrected significance threshold (P < 1.24 × 10^−6^) increased by 10, 14 and 48 to 28, 60 and 55 in the conformation, production, and all groups, respectively (Figure 1E). No significant markers were detected for the reproduction trait category. Across the four trait groups, the metaGWAS^2^ revealed 86 variants, from which 34 were also detected by both other methods, while 21 were detected only by the metaGWAS^1^ (Figure 1B). Including all available samples into the metaGWAS^2^ did not reveal any additional QTL (Figure 2A).

**Figure 2.**
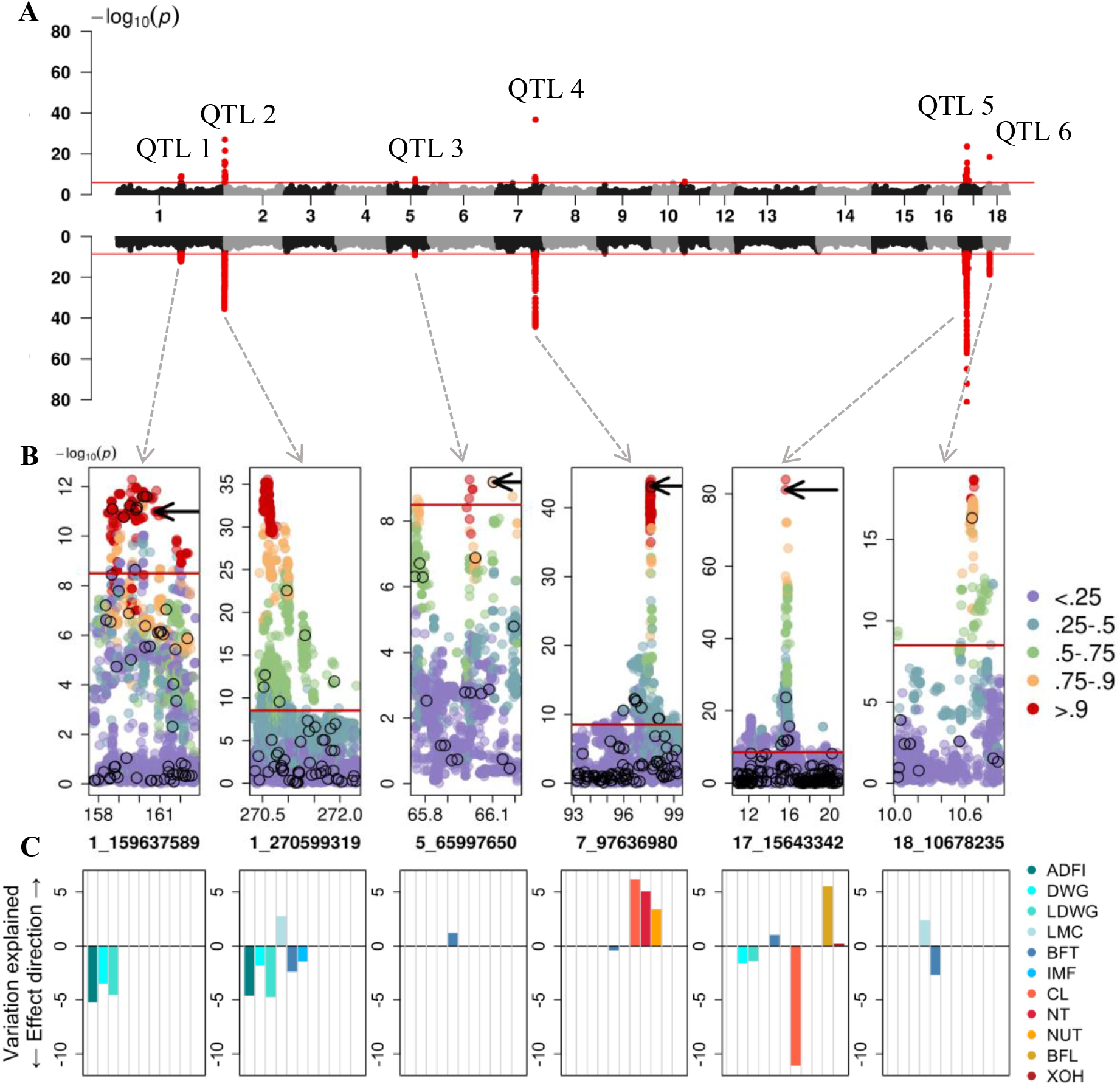
Fine mapping of six QTL detected by metaGWAS. (A) Manhattan plots from array (upper) and imputed sequence (bottom) variants in metaGWAS with 24 traits. (B) Linkage disequilibrium between the lead SNPs and all other variants. Black circles mark array SNPs, arrows point to previously proposed causal variants. The red line indicates the genome-wide Bonferroni-corrected significance threshold. (C) Variation explained (in % of the drEBV variance) by alternative alleles of the lead SNPs in the single traits. Production traits are in blue scale (ADFI - Average daily feed intake; DWG - Daily weight gain on test; LDWG - Lifetime daily weight gain; LMC - Lean meat content; BFT - Back fat thickness; IMF - Intramuscular fat content in loin), and conformation traits in red scale (CL - Carcass length; NT - Number of teats - both sides; NUT - Number of underdeveloped teats; BFL - Bent to pre-bent, front legs; XOH - X- to O-legged).

### Reference panel and imputation of array genotypes to sequence level

Whole-genome sequence-variant genotypes from 421 pigs were used to impute the medium-density genotypes of 5,753 pigs to the whole-genome sequence level. The principal components analysis (PCA; Additional file 9) of a GRM built from 16 million biallelic SNP genotypes confirmed that the sequenced reference panel is representative for our GWAS cohort. Five-fold cross-validation indicated high accuracy of imputation (Additional file 10) with values of 0.92, 0.97 and 0.95 for the squared Pearson’s correlation (R^2^) between true and imputed allele dosages, cross-validated proportion of correctly imputed genotypes (concordance ratio – CR) and model-based accuracies from Beagle5.2 (Beagle DR2), respectively. Although the model-based estimate from Beagle was highly correlated with the Pearson R^2^ (0.81), the Beagle DR2 values were consistently higher.

### Imputed sequence-based association studies

The stGWAS between 24 traits and the 16,051,635 imputed variants revealed 45,288 variants exceeding the Bonferroni-corrected significance threshold in 7 QTL regions which largely agreed with the results from the array-based GWAS (Additional file 11). Two QTL on SSC5 and SSC12 were detected in stGWAS for BFT and NT, respectively, while the other five QTL were associated with multiple traits. The sequence-based GWAS revealed one additional QTL (on SSC12) but did not detect significant association at 5 previously detected QTL likely due to a more stringent Bonferroni-corrected significance threshold resulting from a 350-fold denser marker panel.

A metaGWAS using the summary statistics from stGWAS between the 16,051,635 imputed whole-genome sequence variants and 24 traits using all animals revealed six QTL on SSC1, 5, 7, 17 and 18 (Figure 2, Table 3) with a total of 9,774 variants exceeding the Bonferroni-corrected significance threshold. When the six lead imputed SNPs were fitted as fixed effects in stGWAS, the peaks in the metaGWAS Manhattan plot disappeared (Additional file 12), indicating that the lead SNP accounted for the QTL variance. Four QTL revealed by metaGWAS were significantly associated in stGWAS exclusively with production traits, whereas two QTL (on SSC7 and SSC17) were associated with traits from both the production and conformation categories (Figure 2C). The total trait variance explained per QTL ranged from 0.18 to 11.02 % (Figure 2C).

**Table 3.** Imputed lead variants in pleiotropic QTL revealed by a multi-trait meta-analyses.

| QTL | SSC | Start (bp) | Stop (bp) | Lead SNP | P-value | MAF | Ref \|<br>Alt | #traits | Candidate<br>gene(s) |
| --- | --- | --- | --- | --- | --- | --- | --- | --- | --- |
| **1** | 1 | 157,732,172 | 162,736,434 | 1_159637589 | $5.26 \times 10^{-13}$ | 0.31 | G* \| C | 3 | *MC4R* |
| **2** | 1 | 270,331,332 | 272,315,496 | 1_270599319 | $2.85 \times 10^{-36}$ | 0.14 | C \|<br>CT* | 6 | na |
| 3 | 5 | 65,753,637 | 66,210,538 | 5_65997650 | $5.40 \times 10^{-10}$ | 0.41 | G \| A* | 1 | *CCND2* |
| 4 | 7 | 93,119,538 | 99,261,691 | 7_97636980 | $8.09 \times 10^{-45}$ | 0.34 | A* \| C | 4 | *VRTN*;<br>*ABCD4* |
| 5 | 17 | 10,801,927 | 20,928,904 | 17_15643342 | $1.19 \times 10^{-84}$ | 0.20 | C* \| T | 5 | *BMP2* |
| 6 | 18 | 10,024,756 | 10,904,242 | 18_10678235<br>18_10678293 | $2.00 \times 10^{-19}$ | 0.32 | C \| A*<br>A \| G* | 2 | na |

\* Minor allele in the genotyped dataset; Reference alleles were determined according to the Sscrofa11.1 genome assembly.

#### QTL 1 with lead SNP 1_159637589

A QTL on SSC1 was between 157.73 and 162.74 Mb and encompassed 34,312 imputed sequence variants including 1,157 that were significantly associated in the sequence-based metaGWAS (-log10(P) > 8.5). The QTL was associated with ADFI, DWG, and LDWG, and explained 5.16, 3.45, and 4.49% of the trait variance, respectively. The lead SNP at this QTL was an imputed intergenic sequence variant (rs692827816) located at 159,637,589 bp (P = 5.26 × 10^−13^). rs692827816 was in high LD (R^2^ > 0.90) with 935 variants that had similar P values. One of the variants in high LD was a missense variant (rs81219178) at 160,773,437 bp within the *MC4R* gene, which has previously been proposed as candidate causative variant for growth and fatness traits [35]. rs81219178 segregated at MAF of 0.36 in the SLW population and was imputed from the reference panel with high accuracy (DR2 = 1).

#### QTL 2 with lead SNP 1_270599319

Another QTL on SSC1 was between 270.33 and 272.32 Mb and encompassed 21,683 imputed sequence variants including 3,765 that were significant in the sequence-based metaGWAS (Additional file 13). The QTL was associated with ADFI, DWG, LDWG, LMC, BFT, and IMF, and explained 4.57, 1.77, 4.69, 2.70, 2.35, and 1.40% of the trait variance, respectively. The lead variant was an imputed insertion polymorphism (C>CT) located at 270,599,319 bp (P = 2.85 × 10^−36^), approximately 13 kb downstream from *ASS1* and 53 kb upstream from *FUBP3*. The lead variant was in high LD (R^2^ > 0.90) with 747 variants. The MAF of the lead variant was 0.14 and its genotypes were imputed from the reference panel with high accuracy (DR2 = 0.98).

#### QTL 3 with lead SNP 5_65997650

A QTL on SSC5 was located between 65.75 and 66.21 Mb and encompassed 5,065 imputed sequence variants, from which eight were significantly associated with BFT (Additional file 14). The QTL explained 1.16% of the phenotypic variance of BFT. The lead SNP was an imputed sequence variant located in an intergenic region at 65,997,650 (P = 5.40 × 10^−10^; rs346219461) which was in high LD (R^2^ > 0.90) with seven other variants. rs346219461 was 7.60 kb downstream the fibroblast growth factor 6 (*FGF6*) encoding gene and it had MAF of 0.41, and it was imputed from the reference panel with high accuracy (DR2 = 0.99). The rs346219461 was in LD (R^2^ = 0.82) with a non-coding variant (rs80985094 at 66,103,958 bp, P = 6.41 × 10^−10^) in the third intron of *CCND2*, that was previously proposed as putative causal variant for backfat thickness [34].

#### QTL 4 with lead SNP 7_97636980

A pleiotropic QTL on SSC7 was between 93.12 and 99.26 Mb and it encompassed 31,013 imputed sequence variants including 2,341 that were significant in the sequence-based metaGWAS. The QTL was associated with BFT from the production group (0.37% variance explained), and CL, NT, and NUT from the conformation group, where it explained 6.11, 5.00, and 3.32% of the trait variance, respectively. The lead SNP was an imputed variant (rs333375257 at 97,636,980 bp, P =8.09 × 10^-45^) located 12.7 kb downstream *VRTN*. The rs333375257 had MAF of 0.34 and was imputed from the reference panel with high accuracy (DR2 = 0.99). The rs333375257 was in high LD (R^2^ > 0.90) with 424 sequence variants. A previously described candidate causal variant (rs709317845 at 97,614,602 bp, P = 6.71 × 10^-44^) for the number of thoracic vertebrae [36] was in LD (R^2^ > 0.99) with the rs333375257. In addition, 334 significant variants in the ATP binding cassette subfamily D member 4 (*ABCD4*) gene were detected. This gene was proposed to impact NT in a Duroc population [37] with top SNP rs692640845 at position 97,568,284. In our study, the rs692640845 was highly significantly associated (P = 1.49 × 10^−42^) with CL, NT and NUT, and in almost complete LD (R^2^ = 0.98) with the lead SNP.

#### QTL 5 with lead SNP 17_15643342

A pleiotropic QTL on SSC17 encompassed 82,132 imputed sequence variants including 2,112 that were significantly associated residing between 10.80 and 20.93 Mb. The QTL was associated with DWG, LDWG, and BFT from the production group (explaining 1.56, 1.36, and 0.97% of the trait variance, respectively), and with BL, BFL, and XOH from the conformation group (explaining 11.02, 5.48, 0.18% of the trait variance, respectively). The strongest association was from an imputed sequence variant (rs342044514 at 15,643,342, P = 1.19 × 10^-84^) in an intergenic region 106 kb upstream the *BMP2* gene. The variant had MAF of 0.2 and it was imputed from the reference panel with high accuracy (DR2 = 0.97). The lead SNP was in high LD (R^2^ > 0.90) with two other variants. One of them was a previously proposed candidate causative variant for carcass length [33] (rs320706814 at 15,626,425, P = 8 × 10^−82^) in an intergenic region upstream of the *BMP2* gene, 17 kb away from the lead SNP.

#### QTL 6 with lead SNP 18_10678235

A QTL on *SSC18* between 10.03 and 10.90 Mb encompassed 5,790 imputed sequence variants including 408 that were significant. The QTL was associated with LMC and BFT, explaining 2.34 and 2.62% of the phenotypic variance, respectively. The QTL had two lead SNPs (P = 2 × 10^−19^) in complete LD, which were imputed sequence variants located at 10,678,235 bp (rs338817164) and 10,678,293 bp (rs334203353). Both variants had MAF of 0.32. The imputation accuracy was 0.80. They were in high LD (R^2^ > 0.90) with other 11 intergenic variants.

## Discussion

Single- and multi-trait genome-wide association studies involving array-derived and imputed sequence variant genotypes from 5,753 SLW pigs enabled us to investigate the genetic architecture of 24 complex traits from three trait groups. The response variables for the association tests were deregressed breeding values because the genotyped pigs had progeny-derived phenotypes. Progeny-derived phenotypes have been frequently used to perform association studies in animals that lack own performance records for the traits of interest. To avoid false-positive associations arising from the accumulation of family information in the progeny-derived phenotypes [38], we used the deregressed breeding values and weighed them according to equivalent relatives’ contributions.

The single-trait association analyses revealed 26 trait × QTL associations at eleven QTL of which seven were associated with at least two traits. Exploiting genetic correlations among the traits in a multi-trait framework revealed association for six out of the seven pleiotropic QTL detected in the stGWAS. Despite considering up to 24 phenotypes in the multi-trait association tests, the multi-trait methods applied in our study revealed three QTL, that were not detected by the single-trait association studies. The multi-trait methods did not reveal association at four QTL that were revealed by stGWAS, perhaps because the QTL effects were too small to be detected in our medium-sized cohort.

The mtGWAS and metaGWAS^1^ detected largely the same associated SNPs for almost all trait-groups. However, the metaGWAS^1^ revealed more associated SNPs, and lower P-values for the lead SNPs. Combining all 24 traits in the mtGWAS revealed five associated SNPs, and the metaGWAS^1^ conducted with the same individuals detected association of seven SNPs. Multivariate linear mixed models may suffer from over-parametrisation and loss of power when more than ten traits are considered [39]. The low number of detected QTL might also result from low genetic correlations between the 24 traits, or from a small sample size (N = 1,074 pigs with non-missing records for all 24 traits). The metaGWAS approach enabled us to establish a larger sample size by considering summary statistics from stGWAS that were conducted with a various number of individuals (i.e., some pigs had missing records for some of the traits). In this setting, the number of associated SNPs detected by the metaGWAS^2^ increased to 55 (10-fold higher than before). According to Bolormaa et al. [1], when individuals overlap only partially between the stGWAS, the metaGWAS approach still appropriately considers variances and covariances among the t-values. It is worth mentioning that there are also frameworks that enable considering samples with partially missing phenotypes in multi-trait GWAS [40–42], but these avenues were not explored in the current study.

Our comparisons between GWAS approaches considered microarray-derived SNP genotypes. The imputation of array-derived genotypes up to the sequence level provides more statistical power to identify associated loci because causal variants are in the data and directly tested for association with traits of interest [22, 25]. The pigs in our study were genotyped at 44,733 SNPs. No samples were genotyped with denser (e.g., 600K) arrays precluding the stepwise imputation of genotypes up to the sequence level. The imputation of sequence variant genotypes into sparsely genotyped samples may be inaccurate particularly for rare alleles [24].

However, imputation in our mapping cohort with a haplotype reference panel of 421 sequenced animals was accurate. This is likely because the haplotype reference panel mainly contained animals from the target breed. A meta-analysis of the imputed sequence variant-based stGWAS was conducted to fine-map QTL and prioritise candidate causal variants. We recovered variants that had previously been proposed as candidate causal variants among the top associated variants in four QTL, suggesting that previously reported variants might underpin these QTL also in the SLW breed. However, none of the proposed candidate causal variants was the top variant in our association studies possibly indicating sampling bias [43], presence of multiple trait-associated variants in linkage disequilibrium [44, 45], or that the top variants were inaccurately imputed [46]. It is also possible that the previously reported candidate causal variants are not causal. Further in-depth functional investigations are required to determine and validate the molecular mechanisms underpinning the QTL identified in our study.

Our meta-analyses approach using imputed sequence variants revealed six QTL of which five were associated with multiple traits. Several porcine pleiotropic loci are underpinned by heterozygous loss-of-function alleles that may have fatal consequences in the homozygous state [47–49]. Pleiotropic QTL have also been described in pigs for highly correlated traits [30, 50]. The meta-analyses conducted in our study revealed six pleiotropic QTL including a QTL on SSC17 which is associated with traits belonging to distinct trait categories. This QTL is associated with carcass length (P = 1 × 10^−62^) and daily weight gain (P = 6 × 10^−13^). These two traits are barely correlated with each other (r = 0.02 - 0.05). Another pleiotropic QTL on SSC1 at ~ 270 Mb was associated with six traits ADFI, DWG, LDWG, LMC, BFT, and IMF, that were moderately to highly correlated (mean r ± SD = 0.37 ± 0.27). This chromosomal region harbours QTL for backfat thickness and feed efficiency-related traits in other pig populations [32, 51]. However, candidate causal variants underpinning this QTL had not been proposed so far. The lead SNP in our study was at position 270,599,319 bp, which is 13 kb downstream from the *ASS1* gene. Expression of *ASS1* has been associated with digestive tract development, cell adhesion, response to lipopolysaccharide, and arginine and proline metabolism in pigs [52, 53]. Considering its putative role in energy metabolism, we propose *ASS1* as a positional and functional candidate gene for a pleiotropic QTL at ~ 270 Mb. Further functional annotations of the trait-associated variants in the non-coding regions might help elucidating the genetic mechanism underpinning this pleiotropic QTL.

## Conclusions

Multi-trait associations analyses provide strength of evidence for the presence or absence of a QTL segregating in populations. Here, we compared the multivariate linear model with meta-analyses of single-trait summary statistics using real data. Both approaches performed similarly in correlated groups of traits with complete datasets. The ability of meta-analyses to include different sets of individuals and unrestricted number of traits promoted the detection power. Thus, we recommend using the meta-analyses for getting overview of pleiotropic QTL in cohorts with more than 10 traits. For analyses of reduced and correlated groups of traits, the choice of the method seems to provide indifferent results. Yet, we stress the importance of the single-trait analyses for accurate interpretation of the pleiotropic effects and for the assignment of the affected traits.

The reference-guided imputation to whole-genome sequence level assigned genotypes to 22 million variants with high accuracy. Putative causal variants found in literature were among the top variants in the fine-mapped QTL, supporting the precision of one-step imputation from medium-dense arrays. Our analyses provide overview of the QTL affecting economically important traits in Swiss Large White and might serve as catalogue for future research examining the causal variants for complex traits.

## Methods

### Animals and phenotypes

Deregressed estimated breeding values (drEBV) with their corresponding degrees of determination (r^2^drEBV) and weights (wdrEBV) for 24 traits were provided by the Swiss breeding company SUISAG for 5,753 pigs of the SLW breed. Breeding values were estimated using BLUP multiple trait animal models neglecting genomic information and subsequently deregressed according to Garrick et al. [54]. For all our analyses, we considered drEBV which had r^2^drEBv > 0.3 and were within five standard deviations from the mean values. We considered only traits for which at least 2,000 genotyped animals had records. The final number of animals with phenotypes was between 2,025 for gait (GAIT) and 5,419 for number of teats (NT). Up to 37 % of the pigs had missing records for at least one trait. The traits were assembled in four trait groups (Table 1): reproduction (4 traits), conformation (10 traits), production (10 traits) and all (24 traits).

### Genotypes

Microarray-derived genotypes were available for 17,006 pigs from different breeds. The genotypes were obtained with five SNP panels with medium density. 2,970 pigs were genotyped with the Illumina PorcineSNP60 Bead Chip comprising either 62,163 (v. 1) or 61,565 (v. 2) SNPs; 13,342 pigs were genotyped using customized 60K Bead Chips comprising either 62,549 (v. 1) or 77,119 (v. 2) SNPs; and 546 pigs had genotypes at 68,528 SNPs obtained with the GeneSeek Genomic Profiler (GGP) Porcine 80K array.

We used PLINK (v. 1.9; [55]) to prepare the genotype data. The physical positions of the SNPs were according to the Sscrofa11.1 assembly [56] of the porcine genome. We retained unique autosomal SNPs that did not deviate from Hardy-Weinberg proportions (P < 0.00001), had SNP- and individual-level genotyping rates above 80%, and minor allele frequency (MAF) greater than 0.5%. Sporadically missing genotypes for the resulting 44,733 variants were imputed for 14,292 animals using Beagle (v. 5.0; [57]).

For the array-based GWAS and for the comparison of the multi-trait GWAS methods, we considered 40,382 SNPs that had MAF greater than 5% in 5,753 animals of the SLW breed.

### Genomic heritability

We used the Fisher-scoring algorithm implemented in the GREML module of GCTA (v. 1.92.1; [58]) to estimate variance components while considering the inversed weight of drEBV (wdrEBV). The genomic relationship matrix was built for 14,292 individuals with 44,733 SNPs, but only up to 5,753 SLW animals were used to estimate genomic heritability.

### Imputation to whole-genome sequence level

Whole-genome sequence (WGS) data were available for 421 SLW pigs that had been sequenced at an average read depth of 7.1x, ranging between 2.35x and 37.5x. This panel also included 32 key ancestors of the genotyped SLW pigs that explained a large fraction of the genetic diversity of the current breeding population [16].

Raw sequence data were trimmed and pruned for low-quality bases and reads with default parameter settings of the fastp software (v. 0.20.0; [59]) and subsequently mapped to the Sscrofa11.1 reference genome using the mem-algorithm of the BWA software (v. 0.7.17; [60]). Duplicated reads were marked with the Picard tools software suite (v. 2.25.2; [61]), followed by sorting the alignments by coordinates with Sambamba tool (v. 0.6.6; [62]). The read depth at each genomic position was calculated with the mosdepth software (v. 0.2.2; [63]), considering reads with mapping quality > 10. Variant calling and filtering followed Genome Analysis Toolkit (GATK - v. 4.1.0; [64]) best practice recommendations. Base quality scores were adjusted using the BaseRecalibrator module while considering 63,881,592 unique positions from the porcine dbSNP (v. 150) as known variants. The discovery, genotyping, and filtering of SNPs and INDELs in the 421 pigs was done using the HaplotypeCaller, GenomicsDBImport, GenotypeGVCFs and VariantFiltration modules of the GATK.

The filtered WGS dataset containing 421 pigs and 22,018,148 variants (18,839,630 SNPs and 3,178,518 INDELs) with MAF greater than 0.01 was used for the imputation of sequence variant genotypes into the array dataset. The reference panel was pre-phased using SHAPEIT4 (v. 4.2; [65]) using the --sequencing parameter. The target array dataset was pre-phased using SHAPEIT4 using the phased sequence data as the reference. Sequence variant genotypes were imputed with Beagle (v. 5.2; [57]) with an effective population size of 50. The effective population size was estimated using SneP [66].

The accuracy of imputation was assessed empirically by five-fold cross-validation in the 421 animals as follows: 40 animals which were sequenced at high coverage (>10x), were used as target panel. The remaining 381 animals served as the reference panel. The SNP density in the target panel was reduced to 44,733 SNP chip genotypes and subsequently imputed to the sequence level based on 381 reference animals as described above. The imputed and actual genotypes of the target samples were compared to derive concordance ratio (CR; proportion of correctly imputed genotypes) and squared correlation (R^2^) between imputed and true genotypes.

A genomic relationship matrix (GRM) was built among 14,629 pigs that had 16,387,582 (partially) imputed biallelic SNPs with MAF > 5% (421 reference animals and 14,208 animals with imputed sequence variant genotypes) using GCTA [58]. The first 10 principal components (PC) of the GRM were obtained with PLINK (v. 1.9).

Post-imputation quality control excluded SNPs with MAF < 5%, model-based accuracy of imputation (Beagle DR2) < 0.6, and deviations from Hardy-Weinberg proportions (P < 10^−8^), resulting in a total of 16,051,635 biallelic variants (13,773,179 SNPs and 2,278,456 INDELs) which were used for association analyses and the fine-mapping of QTL in 5,753 SLW pigs.

### Single-trait genome-wide association analysis (stGWAS)

Single marker-based GWAS were conducted between 24 traits (see Table 1 for more information about the traits and the number of individuals with records) and either 40,382 array-derived or 16,051,635 imputed sequence variant genotypes using the mixed model-based approach implemented in the GEMMA software (v.0.98.5; [67]).

The linear mixed model fitted to the data was in the following form: y = W*α* + x*β* + u + *∊*, where y is a vector of phenotypes of *n* animals; W is a matrix of covariates; *α* is a vector of corresponding coefficients; x is a vector of marker genotypes, coded as 0, 1 and 2 for genotype A_1_A_1_, A_1_A_2_ and A_2_A_2_; *β* is the effect of the A_2_ allele; 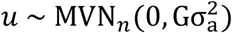 is a random polygenetic effect with G representing the *n* × *n*-dimensional genomic relationship matrix (GRM); 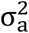 is the additive genetic variance; 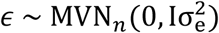 is a vector of errors, with I representing an identity matrix; and 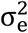 is the residual variance. *MVN_n_* denotes the n-dimensional multivariate normal distribution.

The centred GRM was calculated with GEMMA (--nk 1) using either array-based or imputed sequence variant genotypes. The P-value of each SNP was estimated by the score test implemented in GEMMA (−1mm 3).

The stGWAS was run with either all available individuals or considering only preselected individuals that had non-missing phenotypes within a group of traits. The markers were separately filtered for MAF > 5%. Thus, in the latter run, the stGWAS for the reproduction traits included 41,242 variants typed in 2,553 samples; the stGWAS for the production traits included 40,557 variants typed in 2,689 samples; the stGWAS for the conformation traits included 41,168 variants typed in 1,927 samples, and the stGWAS for the 24 traits together included 41,152 variants typed in 1,074 samples with non-missing records.

### Multi-trait genome-wide association analyses (mtGWAS)

Multi-trait association tests (mtGWAS) were conducted using a multivariate mixed model-based approach implemented in the GEMMA software (v.0.98.5; [39]). The multivariate linear mixed model was parameterised similar to the stGWAS model (see above), except that y, *α*, u, *∊* are matrices with *d* (number of traits) columns, and *β* is a vector with length *d*. 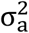 and 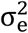 are *d* × *d* symmetric matrices of genetic and environmental variance components, respectively. Because multivariate association testing as implemented in GEMMA requires phenotype data for all individuals and traits, we considered only 2,553, 1,927, 2,689 and 1,074 individuals, respectively, for the reproduction (4 traits), conformation (10 traits), production (10 traits), and all-trait (24 traits) mtGWAS. The GRM, used during the mtGWAS, was the one from the stGWAS.

### Meta-analyses multi-trait genome-wide association (metaGWAS)

A multi-trait meta-analysis (metaGWAS) was conducted with the summary statistics from stGWAS as suggested by Bolormaa et al. [1]. Briefly, the t-values for each marker-trait combination were calculated based on the allele substitution effect and corresponding standard error obtained from the stGWAS. The multi-trait χ^2^ statistic was subsequently calculated based on the *j* × *d* matrix of signed t-values and its *d* × *d* variance-covariance matrix, where *j* is number of markers and *d* is number of traits. P-values for the *j* markers were calculated with pchisq function with *d*-1 degrees of freedom, as implemented in R. We carried out the meta-analyses with the 24 traits classified into the same four trait categories as in mtGWAS (reproduction, production, conformation, and all). First, to enable unbiased comparison between the metaGWAS and the mtGWAS results, we considered summary statistics obtained from stGWAS based on individuals with complete records within the respective trait-group. Second, to increase the power of the association tests through maximizing the volume of entering information, hence exploiting the benefit of the metaGWAS approach, we used the stGWAS summary statistics based on all available individuals for the trait (the first run). For clarity we denote the first and second meta-analyses as metaGWAS^1^ and metaGWAS^2^, respectively. The latter approach was repeated for the fine-mapping of the QTL with imputed sequence variant genotypes.

### Comparison of the association methods

We used a 5 % Bonferroni-corrected significance threshold (1.24 × 10^−6^ and 3.11 × 10^−9^ for array and imputed sequence variant genotypes, respectively) to consider multiple testing. Genomic inflation factors were calculated to compare the distributions of the expected and observed test statistics.

The statistical power was assessed using false discovery rate (FDR). Following Bolormaa et al. [68], the FDR was calculated as 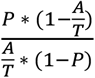, where *P* is the significance threshold (e.g., 1.24 × 10^−6^ or 3.11 × 10^−9^), *A* is the number of significant variants and *T* is the total number of variants tested.

### Fine mapping of detected QTL

We defined QTL as a region of 1 Mb non-overlapping windows, containing at least one significantly associated marker. The marker with the smallest P-value within a QTL was defined as lead variant. Linkage disequilibrium (LD) between the lead variant and all other variants was calculated with the PLINK (v. 1.9) --r2 command. Variants within QTL were annotated with Ensembl’s Variant Effect Predictor (VEP; [69]) tool using local cache files from the Ensembl (release 104) annotation of the porcine genome. The deleteriousness of missense variants was predicted with the SIFT scoring algorithm [70] implemented in VEP.

The proportion of drEBV variance explained by a QTL was estimated with 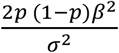, where p is the frequency of the minor allele of the lead SNP and *σ*^2^ is the drEBV variance; *β* is the regression coefficient of the lead SNP. To avoid overestimating the variance explained by a lead variant, we followed the approach described in Kadri et al. [71] and estimated the regression coefficients jointly for all QTL from the stGWAS.

## List of abbreviations

EBV: Estimated breeding value
drEBV: Deregressed estimated breeding value
GWAS: Genome-wide association study
INDEL: Insertion and deletions
LD: Linkage disequilibrium
MAF: Minor allele frequency
metaGWAS: meta-analyses GWAS
mtGWAS: Multi-trait GWAS
PCA: Principal component analysis
QTL: Quantitative trait loci
r^2^drEBV: Reliability of the drEBV
SD: standard deviation
SLW: Swiss Large White
SNP: Single nucleotide polymorph
SSC: *Sus scrofa* chromosome
stGWAS: Single-trait GWAS
wdrEBV: weight of the drEBV
WGS: Whole-genome sequencing

## Declarations

### Ethics approval and consent to participate

No animals were sampled for the present study. Thus, no ethical approval was required.

### Consent for publication

Not applicable.

### Availability of data and materials

High-coverage sequencing read data are available at the European Nucleotide Archive (ENA) (http://www.ebi.ac.uk/ena) of the EMBL at BioProject PRJEB38156 and PRJEB39374.

### Competing interests

AH is employee of SUISAG (the Swiss pig breeding and competence centre). All other authors declare that they have no competing interests.

### Funding

This study was financially supported by SUISAG, Micarna SA and the ETH Zürich Foundation. The funding bodies were not involved in the design of the study and collection, analysis, and interpretation of data and in writing the manuscript.

## Authors’ contributions

Analysis: AN AM HP; Preparation of sequence data: AN ALV HP; Conceived and designed the experiments: AN AM HP; Conceptualisation: HP; Secured funding: HP SN; Wrote the paper: AN HP; Critically revised the manuscript: all authors; Read and approved the final version of the manuscript: all authors.

## Acknowledgements

We thank Claudia Kasper-Völkl for granting access to low-pass sequence data for 294 SLW animals.

## Notes

### Competing Interest Statement

The authors have declared no competing interest.

### Summary of Updates

Updated declarations.

