## Supplementary figures and images for "Comparison of two multi-trait association testing methods and sequence-based fine mapping of six QTL in Swiss Large White pigs" (from another version of this preprint: version 1, 2022-12-15)

### Additional_file1.pdf

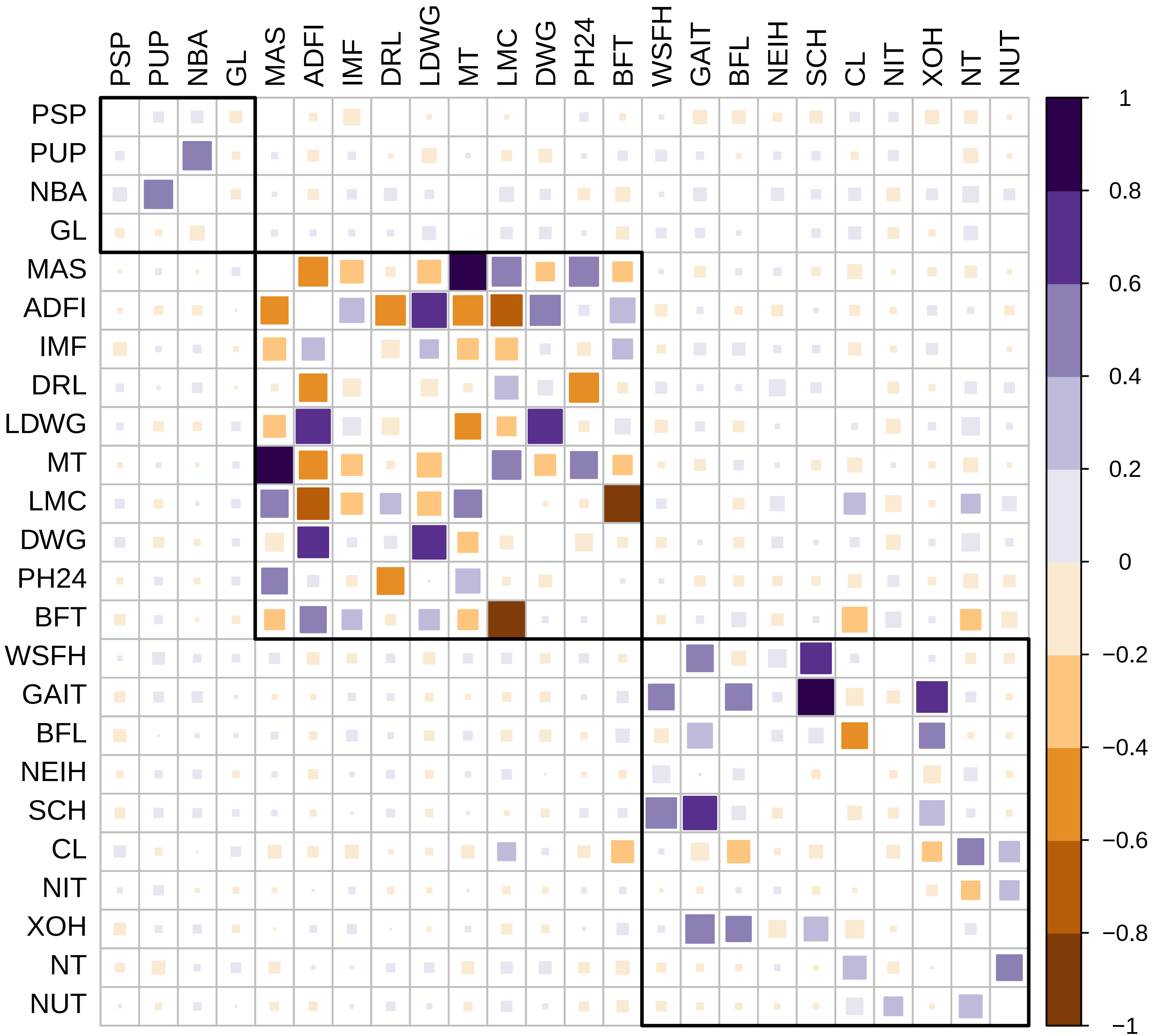

### Additional_file2.pdf

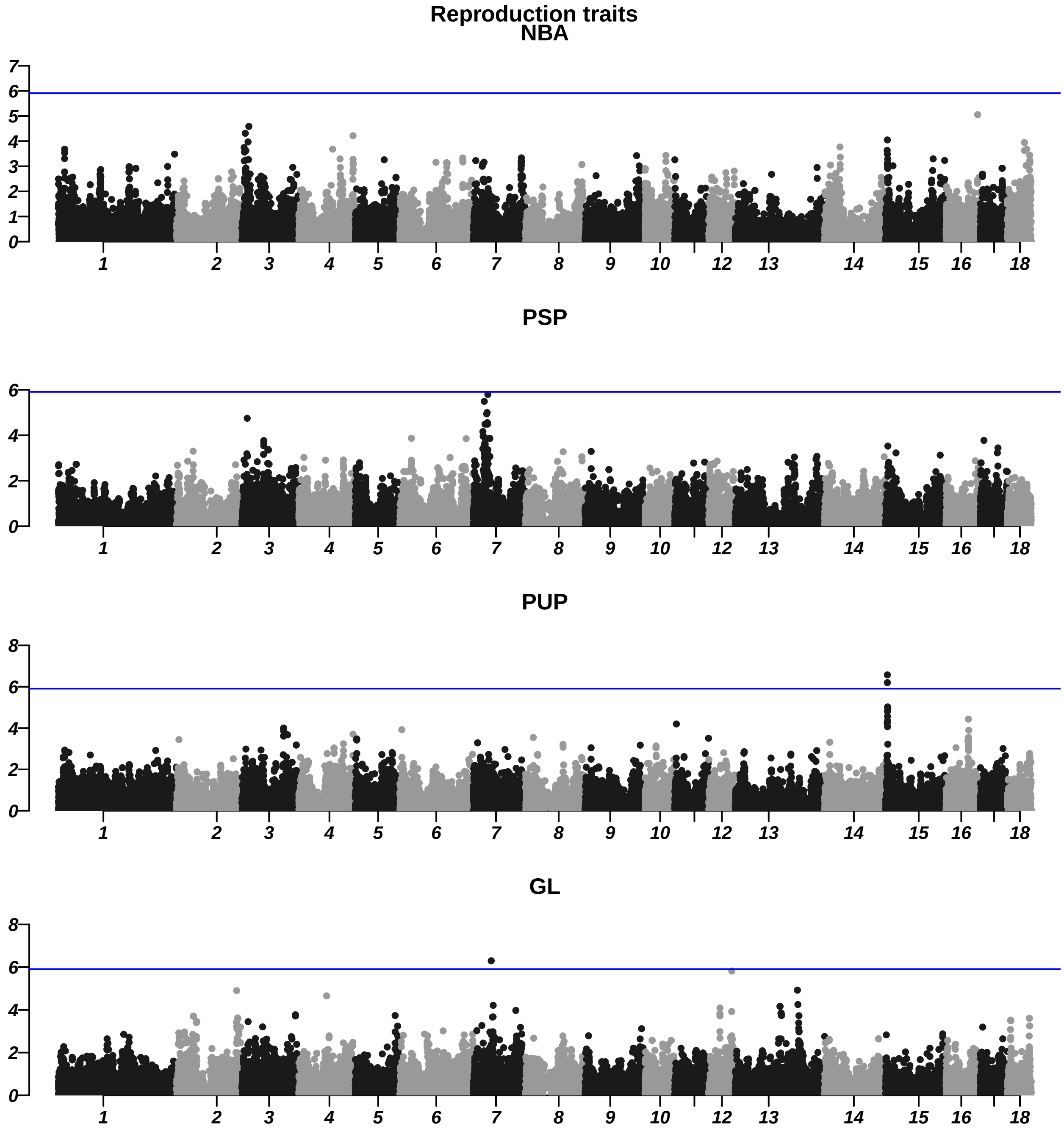

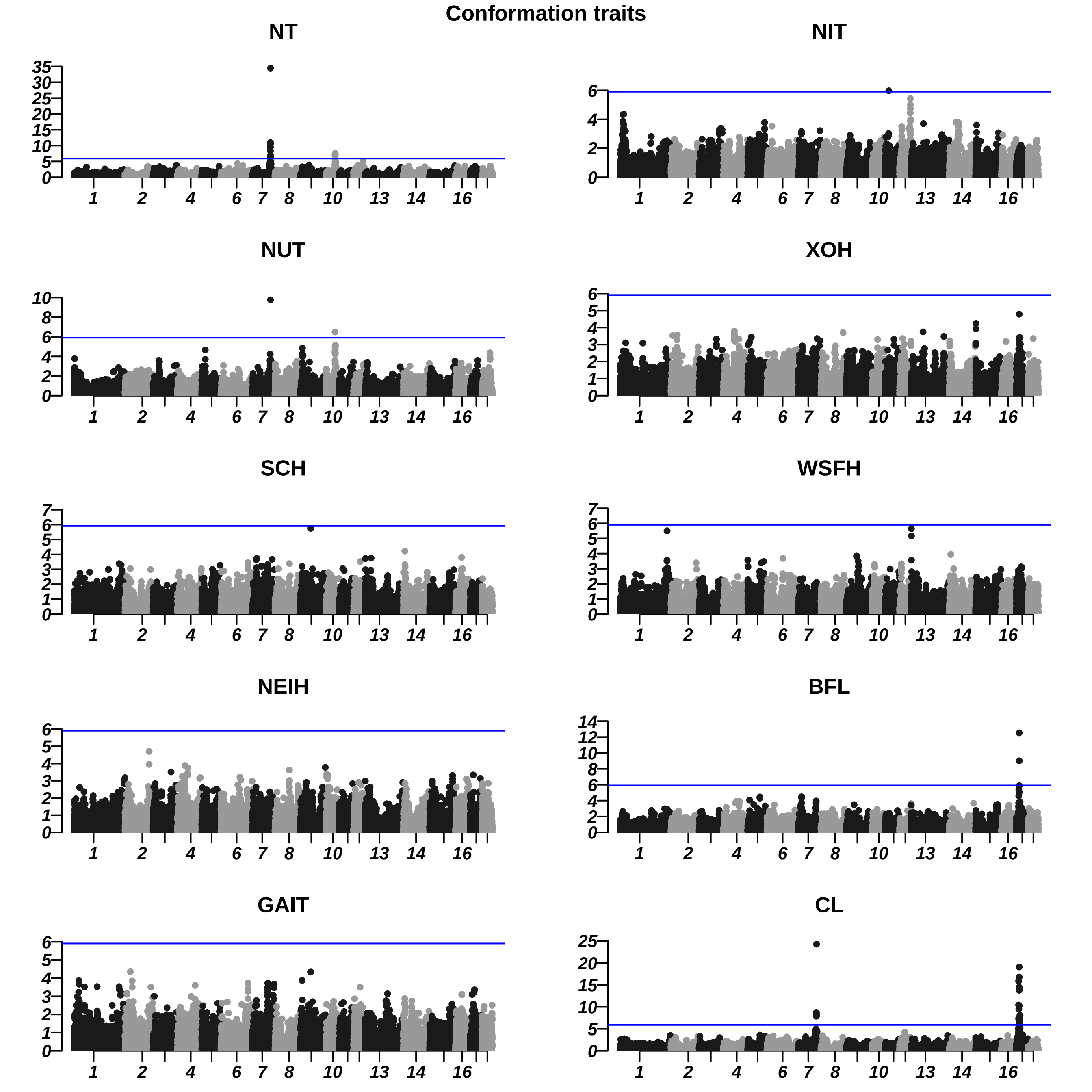

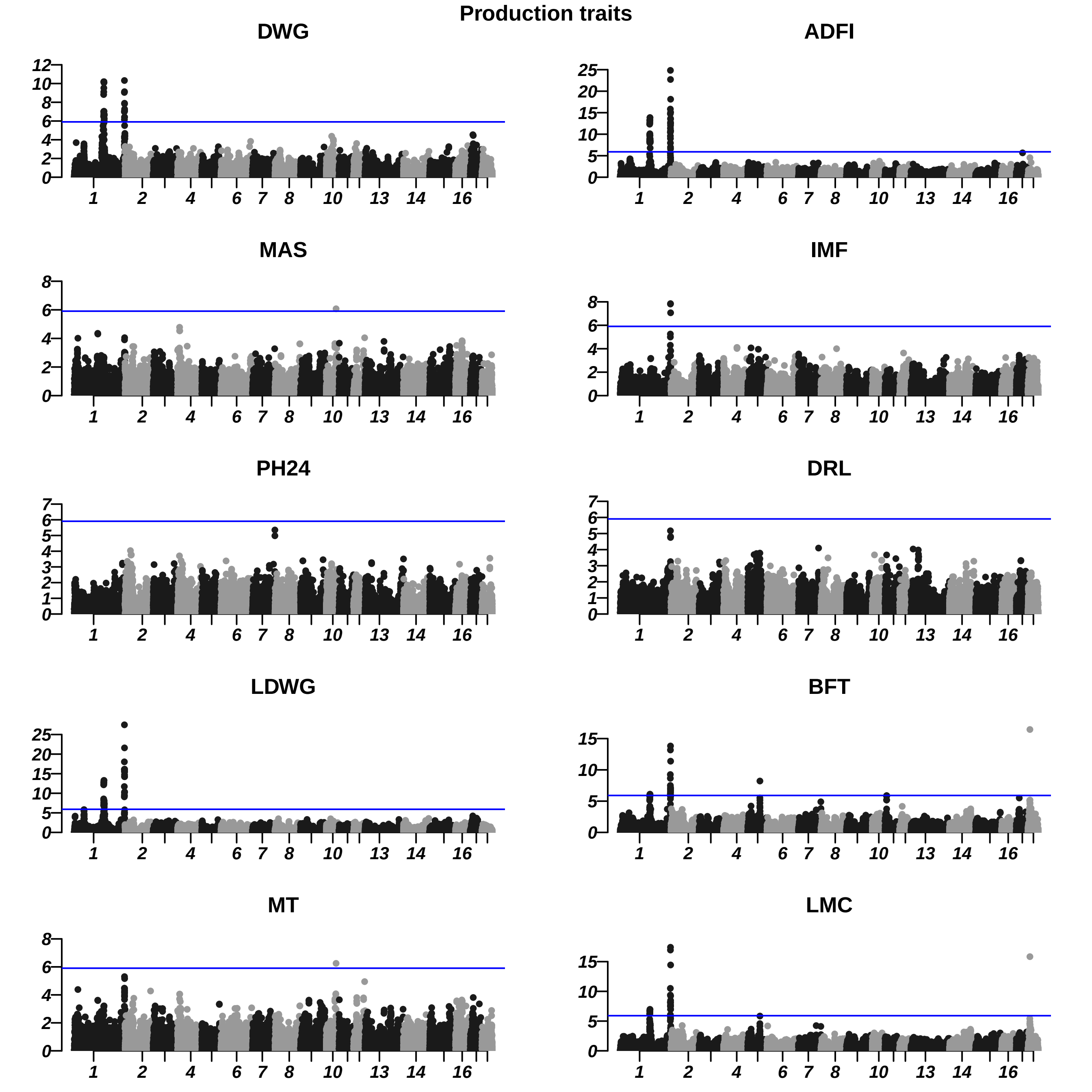

### Additional_file4.pdf

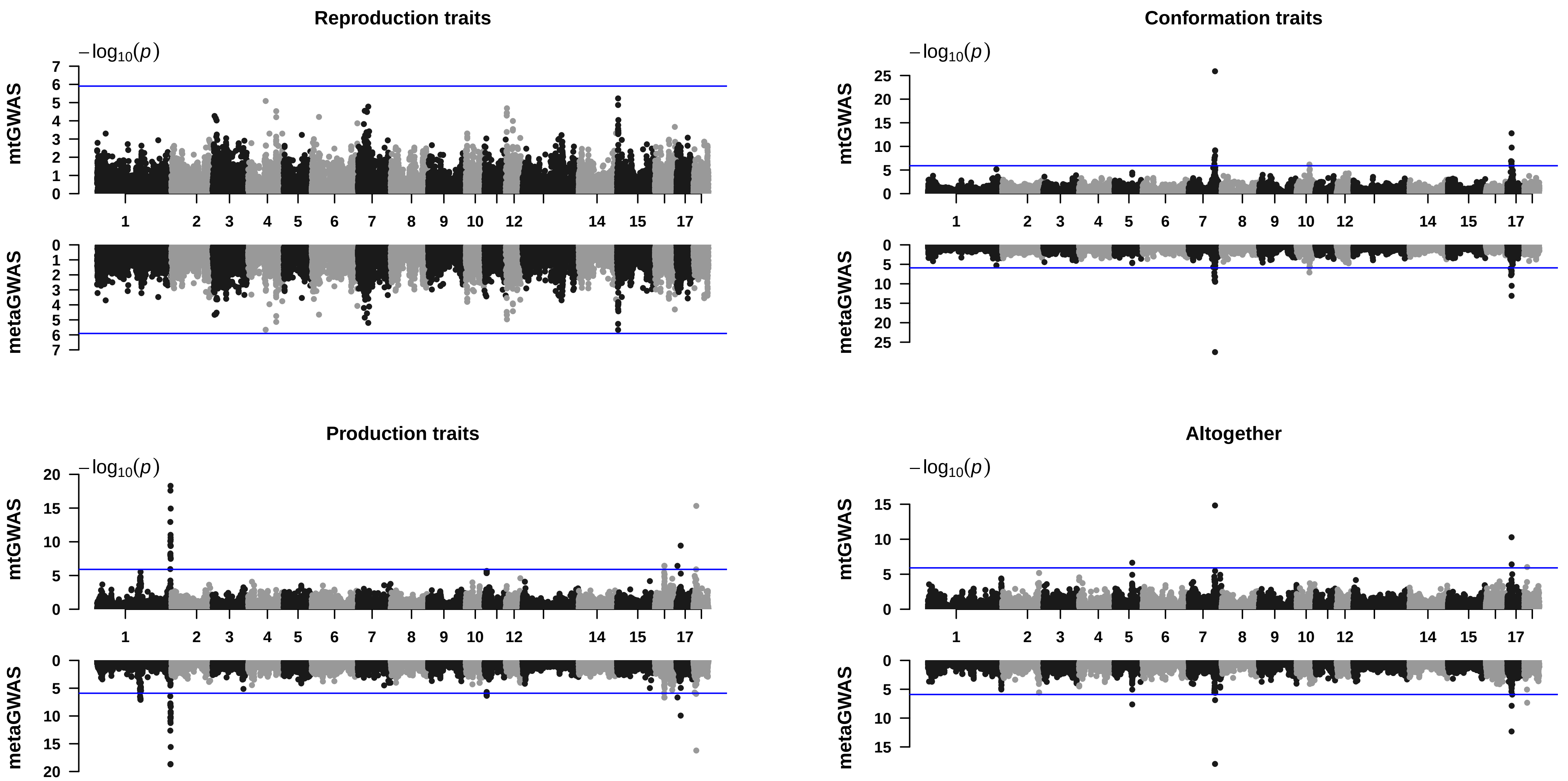

### Additional_file9.pdf

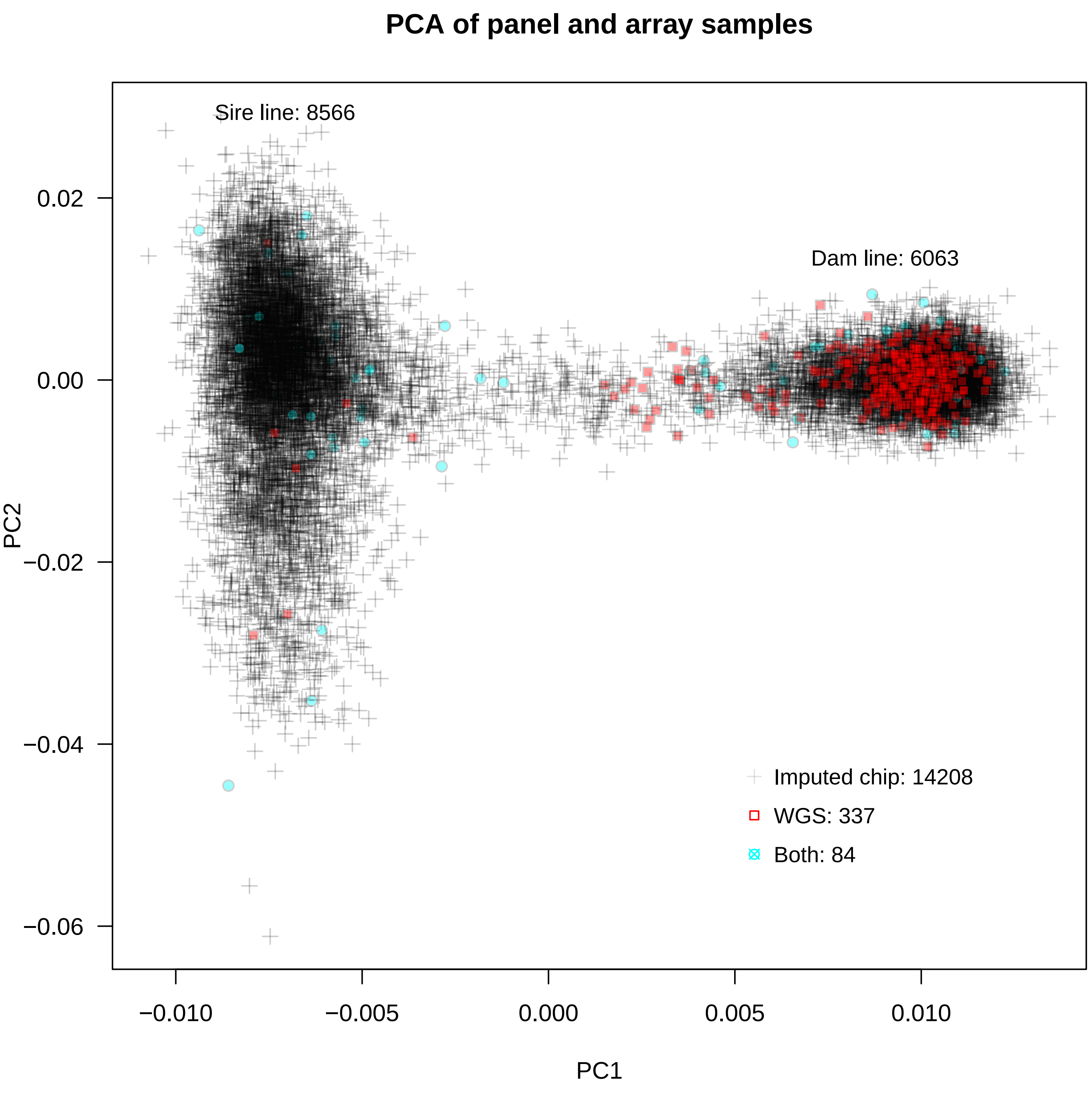

### Additional_file10.png

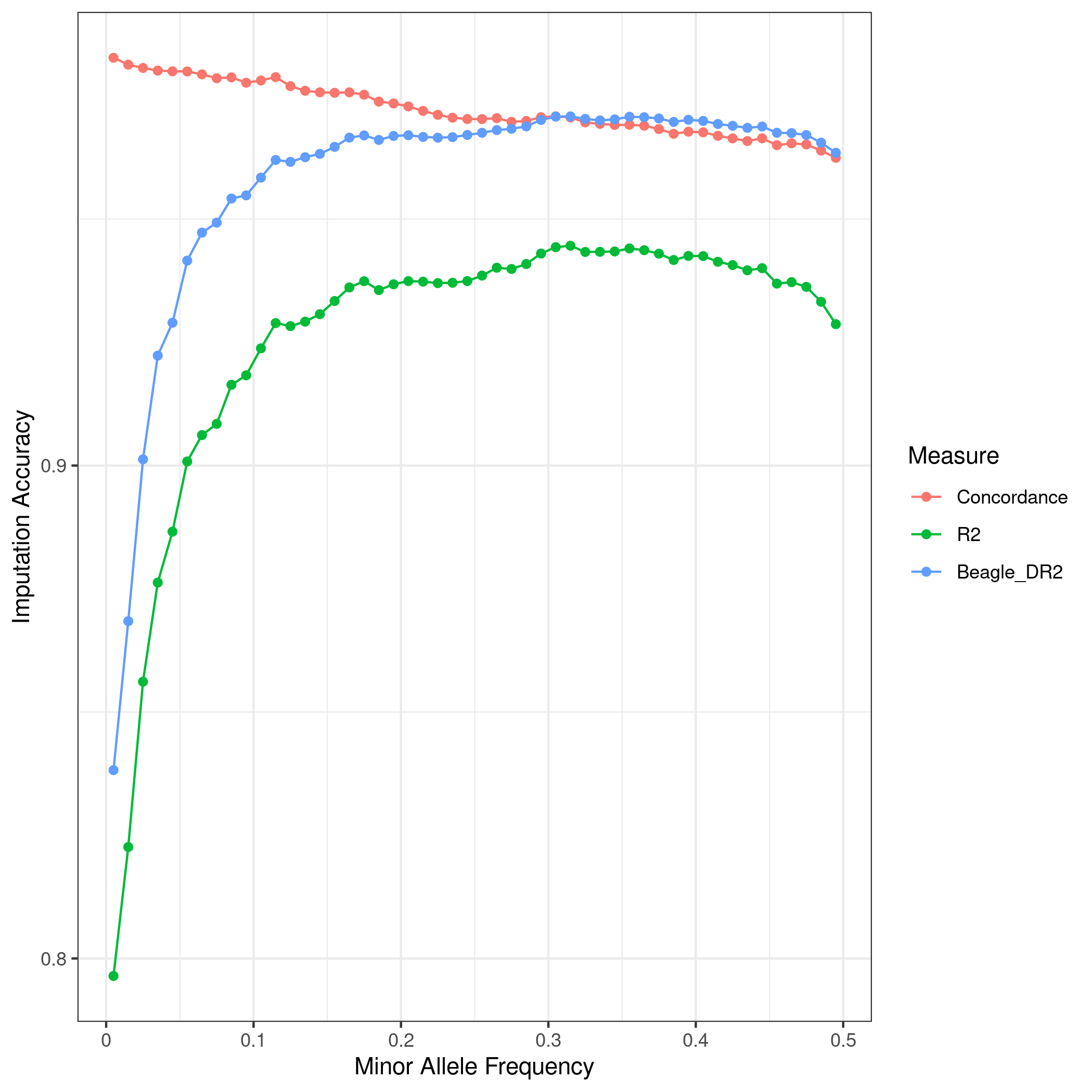

### Additional_file12.tiff

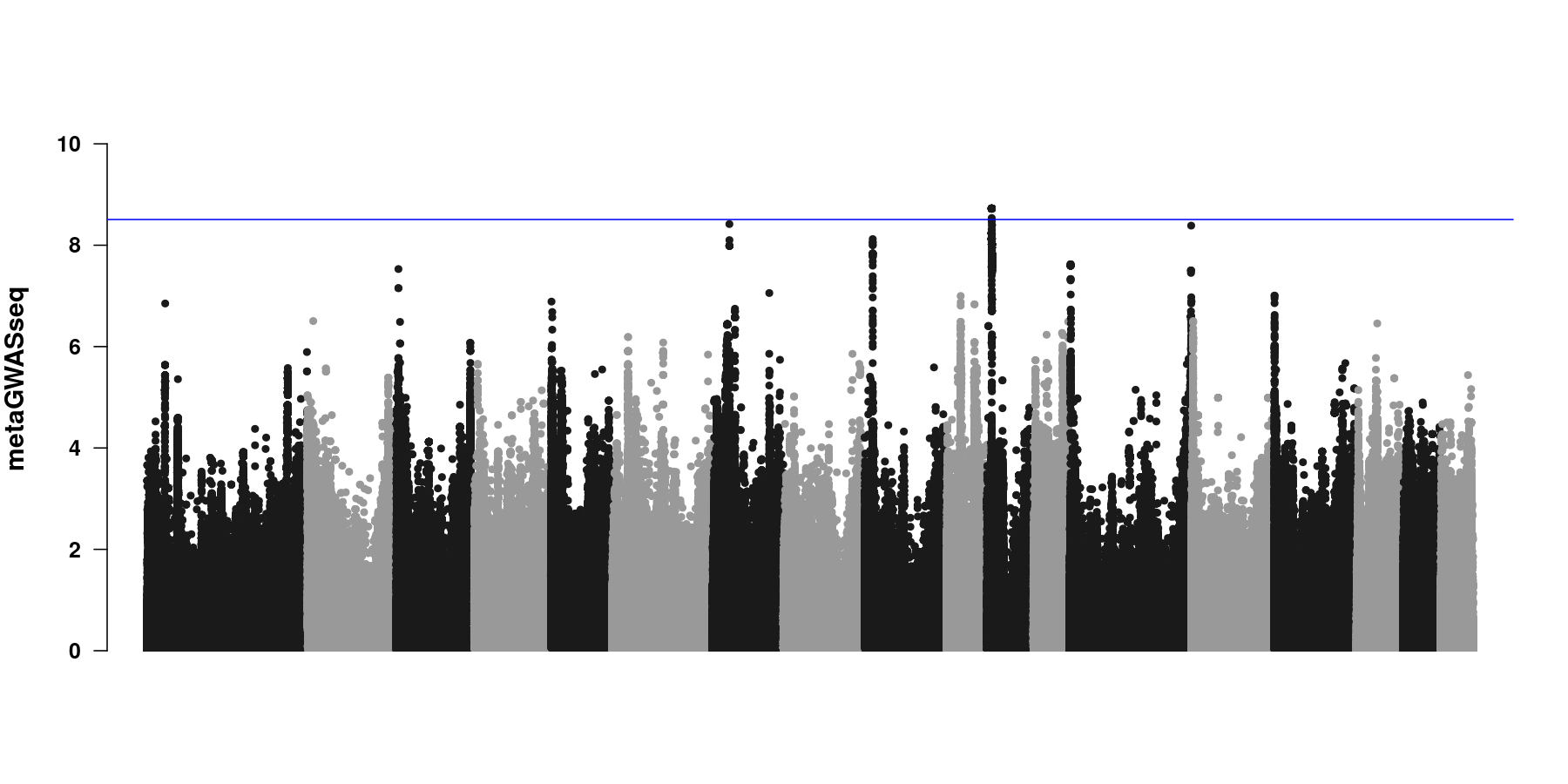
